# Population turnover reverses classic island biogeography predictions in river-like landscapes

**DOI:** 10.1101/170563

**Authors:** Eric Harvey, Isabelle Gounand, Emanuel A. Fronhofer, Florian Altermatt

## Abstract

EH, IG, EAF and FA designed the research; IG and EAF designed the model; IG programmed and ran the model, analyzed the simulation data with support from EAF and produced the figures; EH conducted the lab experiment with support from IG, EAF and FA, processed the experimental data with support from IG, and carried out the analysis of experimental data; all authors participated in results interpretation; EH wrote the first draft of the manuscript; All authors significantly contributed to further manuscript revisions. EH and IG contributed equally to this work.

## Background

Motivated by the global biodiversity crisis, ecologists have spent the last decades identifying biodiversity hotspots. While these global analyses of biodiversity show recurrent patterns of spatial distributions worldwide, unveiling specific drivers remains logistically challenging. Here, we investigate the processes underlying biodiversity patterns in dendritic, river-like landscapes, iconic examples of highly diverse but threatened ecosystems. Combining theory and experiments, we show that the distribution of biodiversity in these landscapes fundamentally depends on how ecological selection is modulated across the landscape: while uniform ecological selection across the network leads to higher diversity in confluences, this pattern can be inverted due to patch size-dependent population turnover. Higher turnover in small headwater patches can slow down ecological selection, increasing local diversity in comparison to large confluences. Our results provide a way forward in the long-standing debate regarding the distribution of diversity in river-like landscapes within a single, experimentally validated conceptual framework.

Local species diversity in dendritic, river-like landscapes is generally expected to be higher in larger, more connected confluences than in headwaters^1–5^. Previous theoretical, comparative and experimental studies have all emphasized the importance of dispersal along the network structure of the landscape as a key driver of this diversity pattern, regardless of the specific dominant local drivers of community dynamics (such as ecological drift^3,6^ or ecological selection^7^). The pattern of lower local diversity in upstream habitats compared to downstream confluences (hereafter ‘classical pattern’), however, is not ubiquitous in natural river systems: recent empirical studies^8,9^ have documented that diversity patterns can be completely reversed, with higher diversity upstream rather than downstream (hereafter ‘reversed pattern’). These contrasting empirical patterns, especially the reversed one, and the transition from one to the other remain insufficiently understood and are currently not accounted for by any theoretical or experimental work. Given that river ecosystems support roughly 10% of all animal species on only 0.01% of the globe surface^10^, it is surprising and disturbing that we are still lacking a general understanding of the processes driving diversity patterns and the transition between seemingly opposed empirical patterns of diversity in these ecosystems. A better understanding of the dominant mechanisms generating contrasting diversity patterns is essential if we hope to counter current trends of erosion of biodiversity and ecosystem services in river ecosystems worldwide^11–13^.

River-like landscapes are structured by spatial flows of organisms and resources. They also inherently display strong environmental gradients, both in terms of resources (habitat size / land use influence) and perturbations (current flow, erosion)^1,14^. While previous theoretical work has primarily focused on dispersal-related mechanisms^3,15,16^, the latter intrinsic characteristics might be key to explaining contrasting diversity patterns in such spatially structured landscapes where habitat size scales with position in the landscape^4,17^: While small headwater patches can be found across a wide range of landscape connectivity, the fewer and larger downstream patches are inherently more connected^14^. The Theory of Island Biogeography^18,19^ (‘TIB’) provides explanation for one of the most established and universal ecological patterns stating that smaller and more isolated patches suffer higher extinction rates and lower immigration rates eventually leading to lower diversity than in larger and more connected patches^18–22^. In such a scenario, competitive exclusion due to ecological selection occurs more rapidly in small headwater communities because, in finite populations, extinction thresholds are reached earlier. Consequently, all else being equals, at equilibrium density, smaller patches should always contain lower diversity. In that context the ‘reversed pattern’ can only occur if (i) all else is not equal and smaller headwater patches contain higher environmental heterogeneity or (ii) equilibrium density is reached more slowly in smaller patches. The former represents the obvious case where headwater patches might contain higher proportion of micro-habitats providing higher niche dimensionality and/or *refugia* from competition and predation. Here we focus on the latter case, where perturbations maintain smaller patches away from their community equilibrium state. This is an especially likely and increasingly widespread scenario in the context of global change^23^. However, due to multiple interacting factors and lack of replication, it is highly challenging to be directly test in the field. To get a causal understanding, we thus develop first a general mathematical model, and then test the main predictions in well-established ecological microcosms.

Perturbations and associated increased population turnover are often seen as a major driver of community dynamics by slowing down competitive exclusion and ultimately favouring species coexistence^24–26^. If perturbations impact smaller patches more strongly than larger patches, it will generate higher population turnover in smaller patches that, in turn, can potentially lead to higher species richness in smaller headwater patches than in larger downstream patches. Thus, in the classical biodiversity pattern, competitive exclusion by ecological selection is expected to occur faster in smaller patches because of finite population sizes, while in the presence of perturbations higher population turnover in smaller patches should slow down competitive exclusion making selection slower in smaller patches thus potentially reversing biodiversity pattern in the landscape.

Indeed, beyond the structural properties of river-like landscapes, specific characteristics of headwaters also suggest that they are naturally more prone to perturbations^27^ and thus to higher population turnover rates. For instance, headwaters exhibit higher benthic surface area to water volume ratio^28^ and thus are more sensitive to minor changes to their surrounding environment^29,30^. Furthermore, they expand into higher elevations, which are experiencing harsher and more variable environmental conditions^31^. Eventually, this asymmetry in population turnover between upstream and downstream patches due to perturbations may generate higher diversity upstream compared to downstream patches.

To resolve the apparent inconsistency in our understanding of local diversity patterns in river-like landscapes^3,8,9,16,32^, we here investigate the processes underlying the distribution of biodiversity and specifically focus on characteristic patch size distributions and population turnover. In natural riverine landscapes, patch size distribution, network configuration and perturbations are intrinsically linked, and cannot be disentangled in a causal approach (see also ^4,14,33^). We thus used a general model reproducing river-like networks properties to disentangle spatial patterns of population turnover from other drivers of biodiversity. We then also validated the predicted turnover-induced reversed biodiversity pattern with a simple experiment. We first show theoretical results from a spatially explicit Lotka-Volterra competition metacommunity model. The model includes dispersal along dendritic networks, demographic stochasticity, and patch size-dependent turnover, that is, higher mortality in smaller patches (see full details in Methods). We secondly corroborated experimentally the model’s main prediction on population turnover effects, using dendritic protist microcosm landscapes^34^ of the same network topologies (see Methods and Fig. 1). Patch size-dependent turnover was experimentally reproduced by sampling, killing and pouring back (no change in biomass) a fixed volume of each community, which resulted in a gradient of population turnover from smaller upstream to larger downstream microcosms (13%, 8%, 4%, and 2% mortality). This mortality gradient implementing an inverse relationship between population turnover and patch size, can be seen as a general effect of environmental perturbation (perturbation-induced turnover).

**Figure 1.**
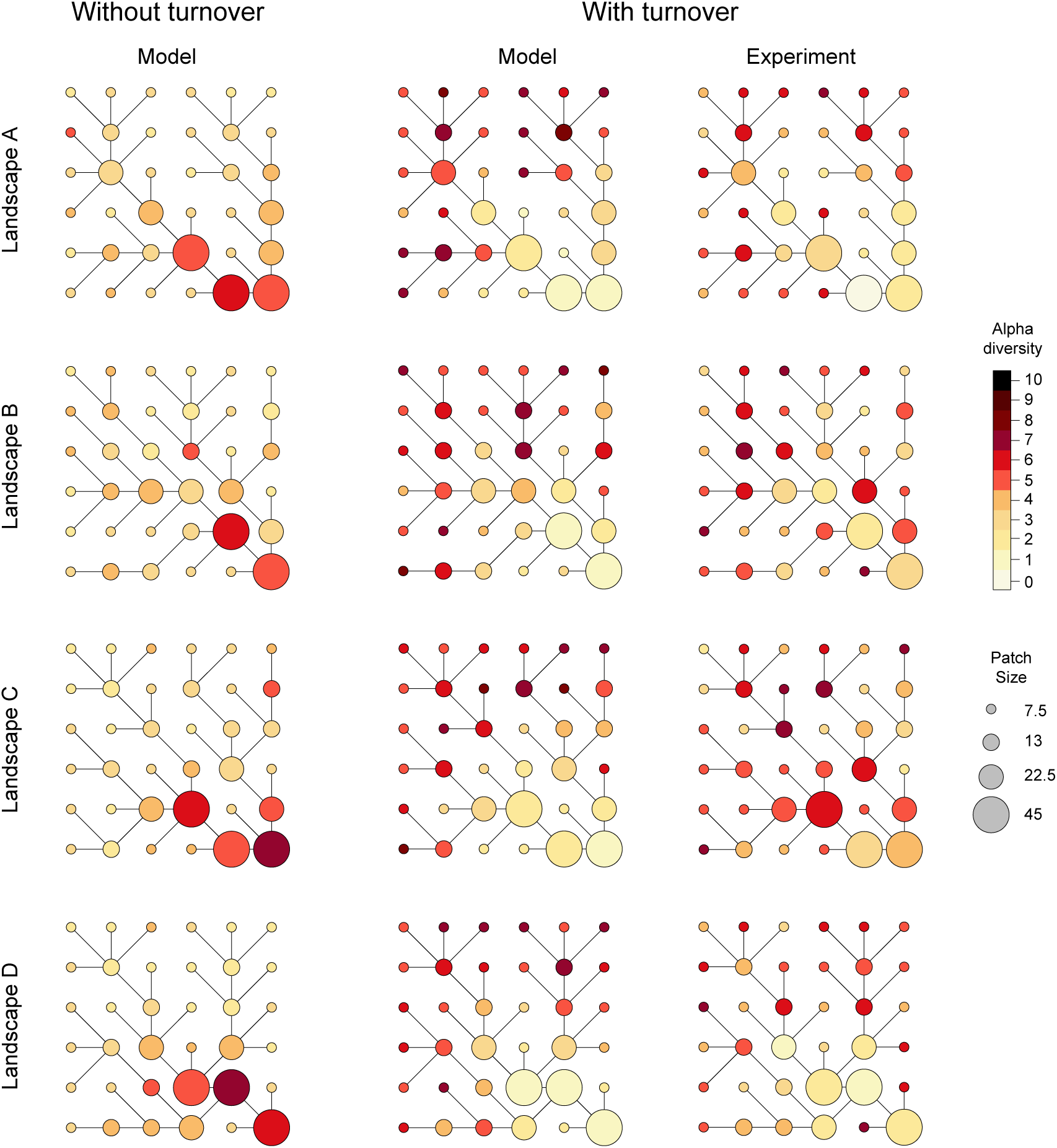
Diversity distribution (local species richness) in dendritic, river-like landscapes with and without patch size-dependent population turnover. The ‘Model’ landscapes displayed in this figure are example of simulations made with the same replicate community. Results were qualitatively similar for each of the 10 replicate communities randomly generated (see Supplementary Figure 1 for results with the different communities; the community used in the present figure is community 7); model parameters specific to these simulations are *d* = 0.05, patch size-dependent mortality rates *m* are 0.1, 0.0864, 0.064 and 0.01 respectively from smaller to larger patches. For network graphs based on experimental results, each graph represents local diversity pattern for one of the four landscape replicates (A, B, C, and D). For illustration purpose, all network graphs are depicted at snapshots of diversity patterns at time steps of comparable local diversity: time steps 118.4 (simulations without population turnover; faster dynamics), 168.1 (simulations with population turnover; slower dynamics), and at experimental day 29 (last sampling day). See complete dynamics over time for model simulations on Supplementary figure 1 and for experimental results on Supplementary figure 2.

## Results

### Classical diversity pattern

In the simulations without perturbation-induced population turnover, the highest diversity levels are found in the large downstream patches (Fig. 1 and 2a). Contrastingly, the smaller, mostly upstream patches have lower diversity levels. This differences in diversity unfolded with smaller patches supporting smaller population sizes (Fig. 3a), which are then more sensitive to stochastic extinctions relative to populations in larger patches (Fig. 3b).

**Figure 2.**
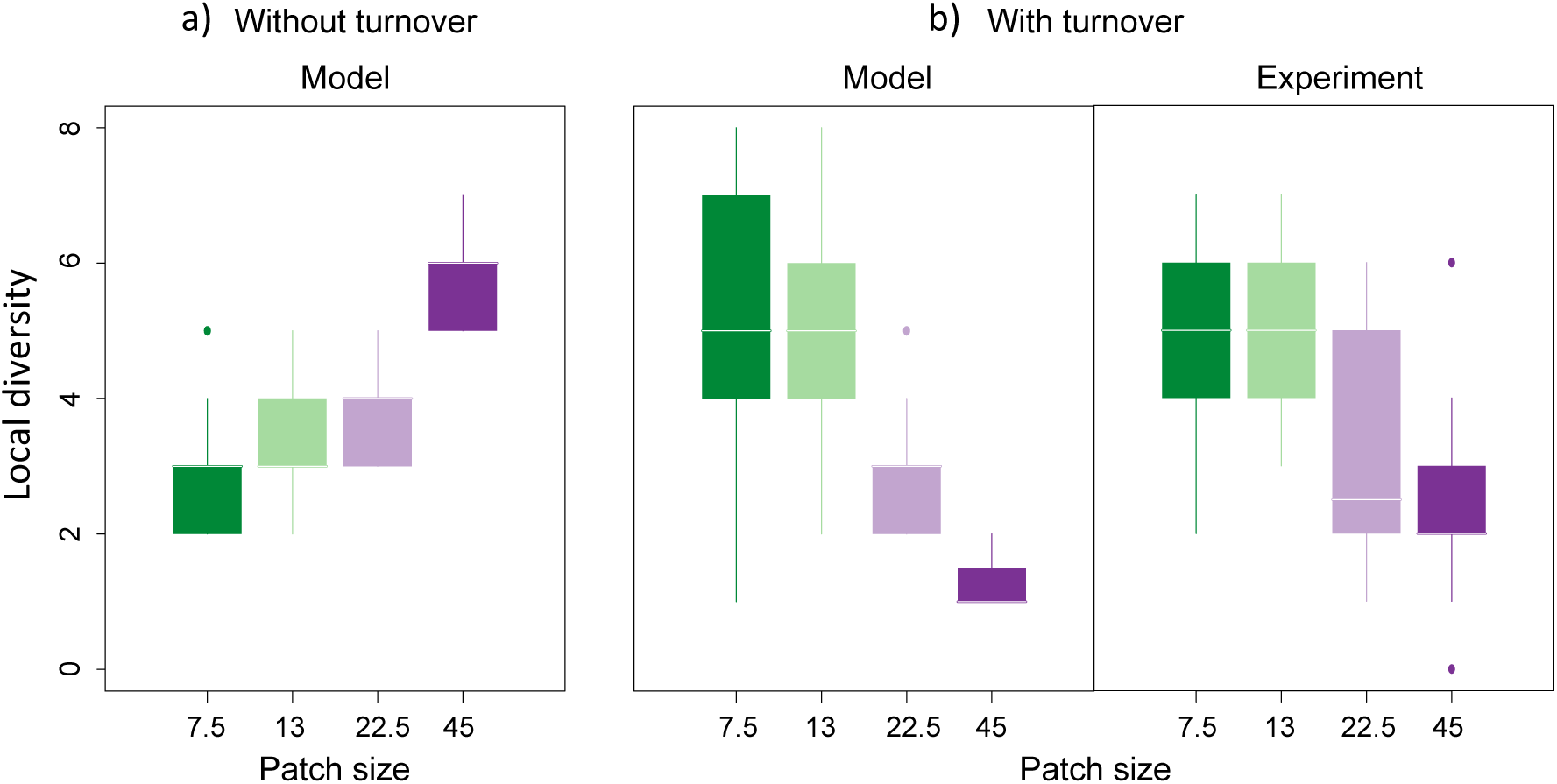
Diversity distribution in dendritic, river-like landscapes a) without and b) with patch size-dependent population turnover. For the panels based on simulations, each boxplot represents the distribution of diversity values across the 4 replicate landscapes used in the experiment, and for one of the 10 replicate communities that were used in the simulations (community 7 in Supplementary figure 1). Parameters and data are the same as in Figure 1. The two largest patch sizes of 45 (N=11) and 22.5 mL (N=14) show significant decline in local diversity compared to the smaller patch sizes of 13 (N=34) and 7.5 mL (N=85) (see Supplementary Table 1 for complete linear mixed effect model results).

**Figure 3.**
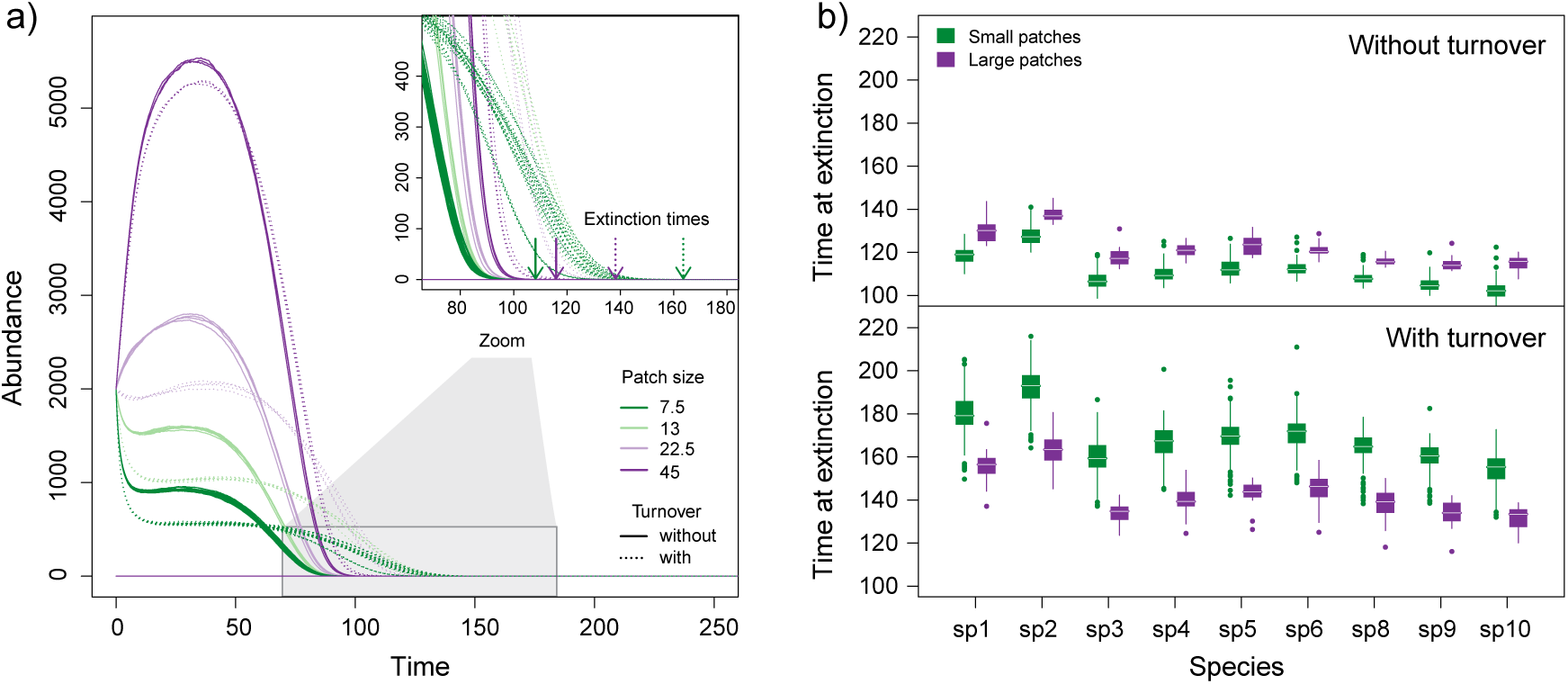
Time to extinction as a function of patch size-dependent turnover in the simulations. a) Temporal dynamics of one species with and without turnover (dotted and full lines respectively). Arrows indicate time at extinction for the smallest (green) and largest (purple) patches with and without turnover. b) Time at extinction for each species in the smallest and the largest patches with turnover (lower panel) or without turnover (upper panel). Each boxplot represents the distribution of values across the 4 replicate landscapes, and for one of the 10 different replicate communities that were used in the simulations (community 7 in Supplementary figure 1). Data and parameters are the same as in Figure 1. Temporal dynamics in a) are the ones of species 10 in community 7 in replicate landscape A.

### Reversed diversity pattern

In the simulations with a gradient of perturbation-induced population turnover, highest diversity levels are found in upstream patches (Fig. 1 and 2b) consistently across all the different community (different species traits) and landscape (different spatial networks) replicates. The same pattern was found in the context of our experiment with an aquatic protist community (Fig. 1 and 2c): As in the simulations (Supplementary Fig. 1), the biodiversity pattern appeared progressively over time with no apparent effects of patch size at the first sampling day (experiment day 7, 7.4 ± 0.67 species in smallest vs. 7.1 ± 0.70 species in largest patches, mean ± sd, Supplementary Fig. 2 and Table 1) compared to the final experimental day 29, when there was a clear and significant decline in local diversity with patch size (4.7 ± 1.3 species in smallest patches vs. 2.5 ± 1.5 species in largest patches, mean ± sd, Fig.1, 2b and Supplementary Table 1). The simulations show that, overall, patch size-dependent population turnover slows down ecological dynamics (ecological selection; Fig. 3a), but does more so in smaller patches (Fig. 3a), which is then increasing the time to extinction of less competitive species (Fig. 3b).

### Persistence of diversity patterns in river-like landscapes

Our model predicts both diversity scenarios to occur within specific parts of the parameter space defined by the magnitude of dispersal and the strength of patch size-dependent population turnover (i.e., slope of the relationship between patch size and population turnover rate, see Methods for details; Fig. 4): weak patch size-dependent population turnover generates the classical diversity pattern, while intermediate to high strengths of patch size-dependent population turnover generate the reversed diversity pattern until population turnover is so high in smaller relative to larger patches that all species upstream go extinct, leading to a return to the classical diversity pattern. As expected, regardless of the strength of patch size-dependent population turnover, increasing dispersal blurs diversity patterns by homogenizing diversity in the landscape (Fig. 4).

**Figure 4.**
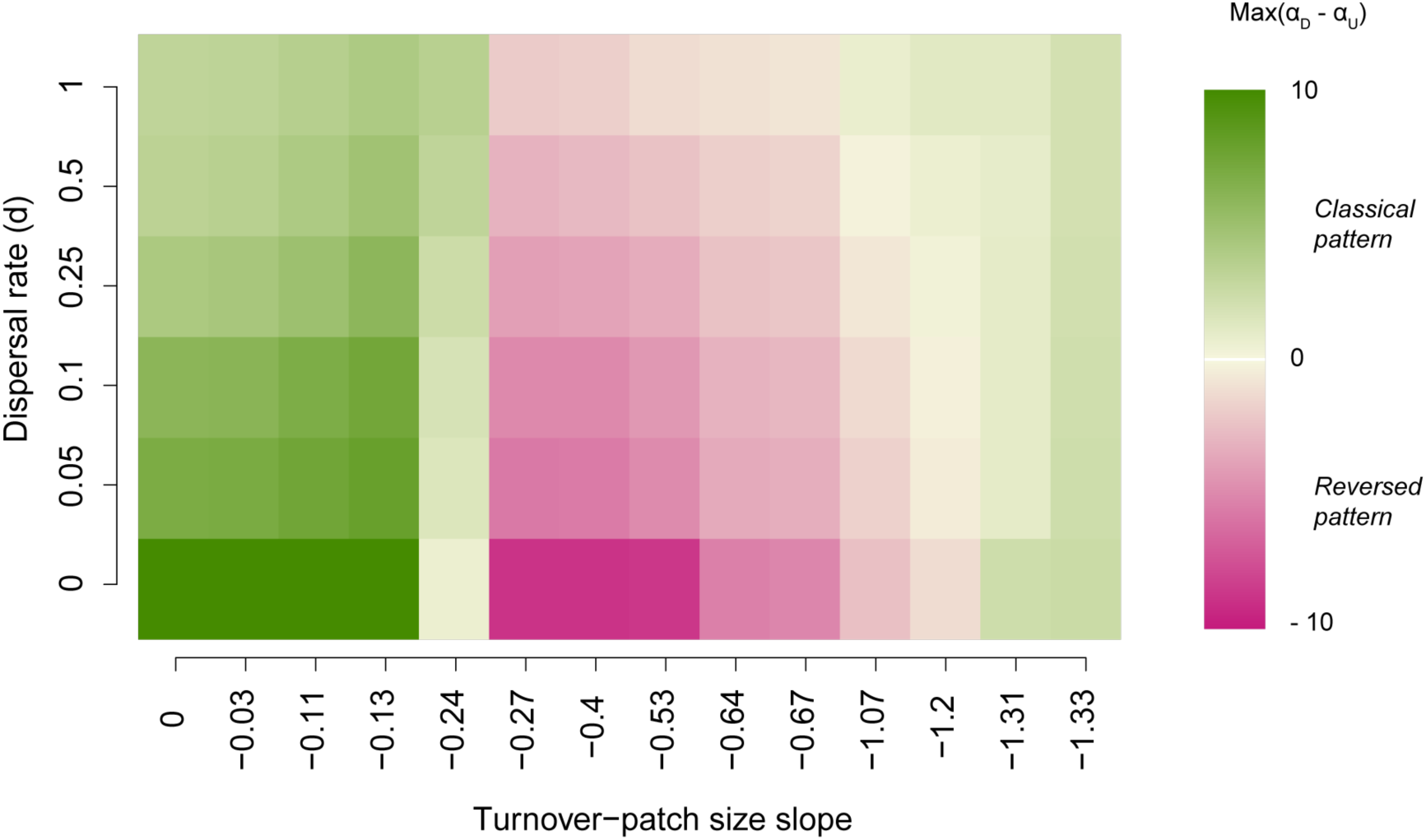
Persistence of diversity patterns in dendritic, river-like landscapes. The figure illustrates the parameter space within which each diversity pattern is found as a function of the strength of patch size-dependent population turnover and the magnitude of dispersal (for all mortality combinations see Supplementary figure 3). α_D_–α_U_ corresponds to the maximum observed difference, along the temporal dynamics, in richness between largest and smallest patches. When α_D_–α_U_ is positive (green colours) the classical pattern emerges over the course of the dynamics, with higher diversity in large than in small patches, while when the difference is negative (pink colours) it is the reversed pattern, with higher diversity in small (i.e., headwarer) rather than in large patches.

## Discussion

Testing the impact of perturbation-induced population turnover on the distribution of biodiversity in river-like landscapes, we found that turnover can alter the classical biodiversity pattern of higher diversity in larger downstream patches predicted by the TIB^3,6,18^: the classical pattern persists only until a certain level of population turnover, at which point asymmetry in turnover between smaller upstream and larger downstream patches will reverse the pattern completely. Such a reversed diversity pattern has been observed in case studies from multiple natural systems^8,9^ and has challenged the idea of one overarching and universal diversity pattern to be expected in river-like landscapes (as proposed by some studies^3,16,35,36^). However, this reversal has hitherto neither been understood mechanistically nor predicted by current theory. Here, we provide a clear mechanistic explanation based on theoretical considerations, which we then successfully validated experimentally.

Dispersal and the landscape context are key factors in driving metacommunity dynamics^37^. Here we showed that specific characteristics of river-like landscapes, namely a combination of the spatial distribution of patch sizes and an associated population turnover gradient, can also play a dominant role in driving diversity patterns. Higher population turnover in smaller headwater patches keeps communities away from their equilibrium state and thus reduces the strength of local ecological selection. This leads to the coexistence of more species for longer periods of time in comparison to larger patches, in which turnover is not strong enough to prevent competitive exclusion. This effect will be especially pronounced in communities with strong competitive asymmetries and even enhanced in systems with competition-colonization trade-offs, as few strong competitors with low dispersal rates, would lead to strong selection in the absence of turnover.

Our work suggests that there may be general mechanisms responsible for the naturally observed diversity patterns including the reversal of diversity patterns. Perturbation-mediated slowing down of competitive exclusion is a well-understood ecological mechanism affecting population dynamics^25,38–40^. Here we show that this mechanism can generate a specific signature in terms of biodiversity patterns when applied over a spatial gradient in perturbation, which are a common feature of river-like landscape. For instance, headwaters in river systems are naturally more prone to higher levels of population turnover because of their higher benthic surface area to water volume ratio compared to larger downstream patches^28^. Headwaters are also more exposed to changes in their surrounding environment, such as natural fire dynamics, potentially leading to more frequent flooding events, higher erosion levels and a decline in terrestrial subsidy (leaf input) on which they are energetically more dependent than larger downstream patches^29,41^. They are on average also located at higher elevation, and thus experiencing stronger variations in temperature and weather conditions. Human-induced perturbations in pristine headwaters are increasingly common^30^ and the general mechanisms demonstrated by our study suggest that these anthropogenic impacts can have unexpected large-scale consequences by inverting biodiversity patterns. This is especially true if perturbations occur in combination with invasive species, which is a common scenario^42^. Thereby, size-dependent population turnover leads to species-poor and highly invasion-prone communities at downstream sections of the river network, whereas species-rich upstream sections may be more resilient due to slower competitive exclusion In conclusion, we show that patch size-dependent turnover, which is intrinsic to river-like landscapes, and its effect on ecological selection can be a key mechanisms responsible for shaping the distribution of biodiversity distribution in river-like landscapes.

## Methods

### Simulation experiment

To investigate the mechanisms underlying biodiversity distribution in dendritic, river-like networks in relation to their intrinsic non-random patch connectivity/size pattern, we first used general simulations of a Lotka-Volterra competition model for 10 species, in which the temporal variation of species *i* density in patch *x*, *N*_*ix*_ is described by:

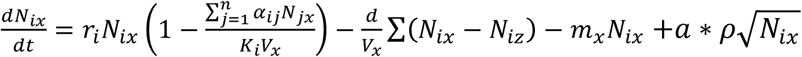

where *r*_*i*_ is the intrinsic growth rate, *α*_*ij*_ the per capita effect of species *j* on species *i* (interaction coefficient), *K*_*i*_ the carrying capacity, which is scaled with the size of the patch *V*_*x*_, *d*_*i*_ the dispersal rate and *m*_*i*_ the patch size-dependent mortality rate (population turnover). In the last term we model demographic stochasticity as in Giometto et al.^43^. At each time step, a proportion *a* times *ρ* of the square root of population density is being either added or removed from each species population in each patch; the parameter *a* modulating the magnitude of this drift is set to 0.25 in all simulations; the random factor *ρ* is sampled for each species within a patch at each time step from a Gaussian distribution of mean *μ* = 0 and variance *σ* = 0.1. Species traits were parameterized in a very simple and general way, independent of our experimental organisms, to preserve the generality of our model predictions. To limit sources of variations, *r*, *K* and *d* are the same for all species in all patches and we implement species differences via interaction coefficients only. Interaction coefficients are drawn from a Gaussian distribution (*μ* = 2, *σ* = 0.25) with all negative coefficients multiplied by -1 to obtain a purely competitive community; *r* and *K* were set to 0.25 and 2000 respectively in all simulations. The dispersal term is a sum function of the population density differences between patch *x* and each patch *z* to which it is connected according to the specific topology of the dendritic landscape. We scale dispersal rates with patch size to make it comparable to the experimental set-up where exchanging a constant volume between two patches was the most feasible dispersal mode. However, the results are robust to change in dispersal mode, using a constant rate either per vertex or per patch. As a minimal assumption to implement patch size dependency in turnover rates, we make mortality rates follow a linear relationship with patch size (see sensitivity analysis below for the values).

For the landscapes, we use river-like networks generated from five different space-filling optimal channel networks^44^ known to reproduce the scaling properties observed in real river systems^4,45^. In order to also be able to use the same landscapes in the experiments, a coarse-graining procedure is used to reduce the 5 generated constructs to equivalent 6x6 patch networks, preserving the characteristics of the original three-dimensional basin (see Fig. 1, for details see appendix A in Carrara et al.^4^), but having a total node size that was experimentally feasible. These 6x6 patch networks contained 36 patches of four different volumes (*V*) respecting river system scaling properties: 7.5, 13, 22.5, and 45 mL (see Fig. 1). These volume values are used for patch sizes *V*_*x*_ in the model.

Using these simulation settings, we test the effect of patch size-dependent population turnover on biodiversity patterns by contrasting simulations with and without turnover. We have two levels of replication: we replicate our simulations over the 5 different dendritic landscapes to assess the independence of the results from specific network topologies, and 10 random realisations of the interaction matrix (replicate communities), leading to 50 simulations total per scenario (turnover settings at a given dispersal rate). Simulations always start with all species already present in all patches and with same absolute density set to 200 individuals. The results are robust to starting conditions with same relative density (scaled to patch size). Simulations are run until 400 time steps, which is sufficient to observe competitive exclusion (see Fig. 3 and Supplements Figure 1 for examples of dynamics). A species for which density falls below 10^−3^ is considered extinct and its density set to 0. The results are robust to the decline or suppressing of this threshold but we keep it to optimize the computation time.

In addition, we explore the persistence of the different observed diversity patterns as a function of the magnitude of dispersal and the strength of the turnover *vs*. patch size relationship (slope of the relationship, see Fig. 4). We explore 6 levels of dispersal with *d* ∈ {0,0.05, 0.1,0.25,0.5,1} and 21 mortality rate combinations between the largest and the smallest patches, that is all combinations with *m* ∈ {0,0.01, 0.05,0.1,0.25,0.5} with mortality in smallest patches being higher or equal to in largest patches (Supplementary figure 3). This results in 14 different slopes of the relationship between turnover (mortality rate) and patch size, ranging from 0 to −1.33. For each parameter combination we run 50 simulations (i.e., 5 replicate landscapes x 10 replicate communities) resulting in a total of 6,300 simulations.

### Experimental validation

To empirically validate the main finding from our simulations, that is, patch size-dependent population turnover leading to a reversal of the ‘classical’ biodiversity pattern in dendritic network, we conducted a protist meta-community experiment. We did not experimentally test for the diversity pattern without perturbations because it has already been empirically verified many times in the literature^2,3,7,27,46,47^, and in the same experimental conditions^4,16^. Therefore, we focussed here on testing specific predictions from the simulations owing to patch-size dependent turnover and the occurrence of the predicted pattern of biodiversity under this scenario. In that context, the experiment constitutes a very conservative test of our much more general simulations. The experiment consisted of seven protist species interacting and dispersing for 29 days along 4 different dendritic networks of 36 patches (same than four of the five landscapes used in the simulations; see Fig. 1). Each landscape had 4 different patch size levels (7.5, 13, 22.5, and 45 mL) connected by dispersal along a dendritic network and preserving the scaling properties observed in real river systems (see Fig. 1 and Supplementary figure 4 for a photo, and section “simulation experiment” above, following Carrara et al.^4^).

Our communities were composed of three bacteria species (*Serratia fonticola*, *Bacillus subtilis* and *Brevibacillus brevis*) fed upon by seven bacterivorous protist and one rotifer species (henceforth called “protists”): *Tetrahymena* sp., *Paramecium caudatum*, *Colpidum striatum*, *Spirostomum* sp., and *Chilomonas* sp., *Blepharisma* sp. and the rotifer *Cephalodella* sp. The latter two species can, next to feeding on bacteria, to a lesser degree also predate on smaller protists. Prior to the beginning of the experiment, each protist species was grown in monoculture in a solution of pre-autoclaved standard protist pellet medium (Carolina Biological Supply, Burlington NC, USA, 0.46 g protist pellets 1 L^−1^ tap water) and 10% bacteria inoculum, until they reached carrying capacity (for methodological details and protocols see Altermatt et al.^34^). Protist abundance and diversity were measured by video recording combined with a trained algorithm to differentiate each species based on their morphological traits (see below).

Each microcosm in our four landscapes consisted of a 50 mL polypropylene falcon tube (VWR, Dietikon, Switzerland). At day 0, we pipetted an equal mixture of each of the seven species into each microcosm to reach the corresponding volume (7.5, 13, 22.5 or 45 mL). Thus, protist communities were added at 15% of their carrying capacity and were allowed to grow 24 hours before the first dispersal event. Dispersal and imposed population turnover (see below) occurred two times per week, while sampling of the communities for species count was done once a week (two dispersal events between each sampling with always at least 48 hours between the last dispersal/turnover event and sampling). Sampling events and counting were done at day 0, 7, 15, 21, 29 of the experiment, while dispersal and turnover events occurred at day 1, 4, 8, 11, 16, 19, 22, 25 of the experiment.

Dispersal was done by pipetting a fixed volume per vertex (1 mL) to each of the connected patches. Dispersal was bi-directional along each vertex (1 mL from a to b and 1 mL from b to a), which ensured the maintenance of the same volume in each patch throughout the duration of the experiment. Other experiments in the same settings and simulations have shown that directional dispersal tends to strengthen meta-population and community patterns compared with bi-directional dispersal, but with no qualitative differences in the observed patterns^4,16,48^. Thus, doing dispersal in this way is conservative. For dispersal we used a mirror landscape (following methods developed in Carrara et al.^16^): first, 1 mL was sampled for each vertex connecting a microcosm in the real landscape and then pipetted to the recipient microcosm, but in the mirror landscape. Once this was done for all microcosms, the content of each mirror microcosm was poured to the same microcosm in the real landscape. It is noteworthy, however, that because we started our simulations and experiment with species already assembled (all species were present at start in all patches), we probably are underestimating the importance of dispersal dynamics in the process of early assembly^49^.

Patch size-dependent population turnover was experimentally reproduced by sampling, killing and pouring back (no change in biomass) a fixed volume of each community, which resulted in a gradient of turnover from smaller upstream to larger downstream microcosms (13%, 8%, 4%, and 2%), and can be seen as the most general type of disturbance. For each mortality event, 1 mL was sampled from each microcosm and microwaved until boiling to turn all living cells into detritus^50^. After a 1-hour cooling period at ambient temperature (20 °C), the microwaved sampled had reached ambient temperature again and were poured back into the same microcosm.

At each measurement day, 0.4 mL was sampled from each microcosm for the protist density measurements. Protist density was measured by using a standardized video recording and analysis procedure^51,52^ (for details see Supplementary appendix A). In short, a constant volume (34.4 μL) of each 0.4 mL sample was measured under a dissecting microscope connected to a camera for the recording of videos (5 s per video, see Supplementary appendix A for further details on this method). Then, using a customized version of the R-package bemovi^52^, we used an image processing software (ImageJ, National Institute of Health, USA) to extract the number of moving organisms per video frame along with a suite of different traits for each occurrence (e.g., speed, shape, size) that could then be used to filter out background movement noise (e.g., particles from the medium) and to identify species in a mixture (see Supplementary appendix A).

### Statistical analysis

To test for the effect of patch size on protist local diversity we used a two-way linear mixed effect model testing the interactive effects of patch size and continuous time on protist species richness. To control for temporal pseudo-replication, we added replicate and time as nested random factors. The model was fitted by maximizing the restricted log-likelihood (“REML”, see ^53^). The linear mixed effect model was conducted using the R-package NLME^53^. The complete results can be found in Supplementary Table 1.

### Data availability

The main data and r-script to reproduce the experimental results can be downloaded from Dryad (DOI:XX.XXXX/Dryad.XXXXXX).

## Acknowledgement

We thank S. Gut, S. Flückiger and E. Keller for help during the laboratory work. Funding is from the Swiss National Science Foundation Grant PP00P3_150698

